# Chronic ER Stress Disrupts Mitochondrial-Associated ER Membrane Integrity in Corneal Endothelial Cells

**DOI:** 10.64898/2026.01.09.698664

**Authors:** Stephanie Lee, Stefan Y. Kim, H Orhan Akman, Saba Qureshi, Anisha Kasi, William Steidl, Lukas Ritzer, Mariane Price, Francis W Price, Eric A. Schon, Varun Kumar

## Abstract

**Purpose:** To characterize alterations in MAMs and identify key MAM proteins associated with ER and mitochondrial stress in degenerating corneal endothelium, as seen in diseases like Fuchs’ endothelial corneal dystrophy (FECD).

**Design:** Experimental study.

**Subjects, Participants, and/or Controls:** A total of 5 FECD human donor corneal endothelium were obtained from Massachusetts Eye and Ear and from Price Vision Group in the US for TEM imaging. Donor corneas without known ocular pathologies were purchased from SightLife (Pennsylvania, US) for use as controls against diseases tissue samples.

**Methods:** Human corneal endothelial cell line (HCEnC-21T) and Fuchs’ corneal endothelial cell line (F35T) were cultured and subjected to ER stressor tunicamycin (1, 10 μg/ml) for 6, 24, or 48 hours. MAM proteins were isolated by subcellular fractionation, where key ER and mitochondrial damage sensor proteins were identified by immunoblotting. Functional MAM protein expression was assessed through immunofluorescence staining. ER-mitochondrial contact sites were quantified through immunofluorescent detection of transfected MAM plasmids in normal and Fuchs’ cell lines. ER and mitochondrial interactions were analyzed in 21T and F35T, alongside normal and Fuchs’ human tissue samples, using transmission electron microscopy (TEM).

**Main Outcome Measures:** Detection of contact distance disruption between ER and mitochondria following ER stress. Identification of MAM protein alterations in stressed corneal endothelium.

**Results:** Contact distance between ER and mitochondria was significantly increased in Fuchs’ tissues, with a similar finding observed in 21T cells following tunicamycin treatment. Subjugation of 21T HCEnCs to tunicamycin for 6 hours saw an elevation in fluorescent intensity of MAM plasmids, however, prolonged treatment with ER stressor at 48 hours resulted in marked reduction of MAM plasmid intensity. Fluorescent intensity quantification of phosphatidylserine synthase 1 (PTDSS1), a functional MAM protein, in 21T cells showed lowered expression after 24 hours of tunicamycin treatment. ER stress protein (PERK) and mitochondria damage sensor protein (Parkin) expressions were significantly increased in MAM fractions 24 hours post tunicamycin treatment in HCEnC-21T.

**Conclusions:** Reduced ER-mitochondrial communication, alongside enrichment of mitochondria and ER stress proteins in the MAM of HCEnCs under chronic ER stress, suggests the importance of MAMs as potential mediators of ER-mitochondrial crosstalk, highlighting the significant role they may play in facilitating corneal endothelial degeneration.

## Introduction

Fuchs’ endothelial corneal dystrophy (FECD) is a bilateral, genetically heterogeneous,^1^ age-related, female-predominant,^2^ degenerative disorder^3^ of the corneal endothelium. The current estimated global prevalence of FECD is 300 million people aged 40 or older, with an expected rise to 415 million affected by 2050 ^4^ FECD is characterized by the formation of extracellular matrix (ECM) deposits, known as guttae, on the Descemet’s membrane, leading to corneal edema and progressive loss of corneal endothelial cells (CEnCs) ^5^ Currently, the only treatment for FECD is corneal transplantation, a costly and invasive option, thus underscoring the need for more pharmacological treatments.

Two major factors contributing to CEnC apoptosis in FECD are endoplasmic reticulum (ER) and mitochondrial stress. ER stress is known to activate the unfolded protein response (UPR),^6^ comprising of three signaling pathways: protein kinase RNA-like ER kinase (PERK; gene *EIF2AK3*), inositol-requiring protein 1 (IRE1), and activating transcription factor 6 (ATF6). In FECD, it has been postulated that excessive synthesis of ECM proteins leads to their abnormal accumulation in the corneal endothelium, contributing to the formation of guttae, possibly activating ER stress pathways of the UPR, and ultimately leading to CEnC apoptosis.^7^ Mitochondrial stress is another important factor in FECD. Stimuli, such as oxidative stress, can trigger the mitochondria-mediated intrinsic apoptotic pathway,^8^ disrupting mitochondrial membrane potential, releasing cytochrome C, and eventually resulting in cellular apoptosis.^9^ Furthermore, unresolved mitochondrial stress can lead to elevated mitochondrial fragmentation, contributing to increased mitophagy. Subsequently, this result in the reduction of mitochondrial mass,^10^ possibly fueling CEnC loss in FECD due to the inability to meet energy demands of the cells. Although ER and mitochondrial stress have been independently associated with the pathogenesis of Fuchs’ endothelial corneal dystrophy, there have been limited studies investigating their crosstalk in FECD. Our lab previously observed the activation of all three ER stress pathways following ER stress induction in human corneal endothelial cells through exposure to tunicamycin, supporting the assertion that ER stress contributes to corneal degradation like FECD by activating UPR.^11^ Interestingly, we also found that treatment of 21T HCEnCs with tunicamycin led to changes in mitochondrial bioenergetics and dynamics, including the loss of mitochondrial membrane potential, decreased ATP production, mitochondrial fragmentation, and to the induction of the mitochondrial-mediated intrinsic apoptotic pathway, demonstrating the presence of crosstalk between ER and mitochondria in degenerative corneal endothelial conditions.^11^

ER and mitochondrial communication via mitochondria-associated ER membranes (MAMs) has been established in other disorders, such as Alzheimer’s disease and Parkinson’s disease.^12^ MAMs are sites of physical contact and communication between ER and mitochondria, and are involved in a variety of cellular processes, including calcium signaling, lipid synthesis and transfer, ER stress, and mitochondrial dynamics.^13^ Specifically, prior studies have found enrichment of ER stress sensor proteins from the UPR pathway, RNA-dependent protein kinase-like ER kinase (PERK), in the MAM of murine embryonic fibroblasts (MEF), and its deficiency resulted in decreased ER-mitochondria contact sites, with loss of PERK expression significantly disrupting calcium signaling in MEFs subjected to ER stress.^14^ Reintroduction of PERK saw the restoration of reactive oxygen species (ROS) signal propagation from ER to mitochondria in ROS-mediated ER stress-treated cells, underscoring the role of ER UPR proteins in facilitating physical and functional communication between the ER and mitochondria.^14^ Mitochondrial damage sensor protein Parkin, a 465-amino-acid E3 ubiquitin ligase responding to mitochondrial membrane potential (MMP) loss, is another similarly relevant player in this MAM space.^15^ Parkin activates proteasomal degradation via polyubiquitination of various mitochondrial proteins and mediates mitophagy, translocating to MAMs and aids in regulation of ER-mitochondrial contact sites through mono-ubiquitination. ^16,17^

Given our lab’s previous finding of ER-mitochondria crosstalk in FECD, in combination with existing literature identifying MAMs as sites of ER-mitochondria communication, we sought to explore mitochondrial-associated ER membranes as potential target sites for ER-mitochondria crosstalk in corneal endothelium and aim to elucidate the effects of ER stress on MAMs in CEnCs. We hypothesize under chronic ER stress, mitochondria-associated ER membranes (MAM) and their contacts are disrupted, affecting normal subcellular organization of ER and mitochondria as well as regular functions of MAMs in CEnCs. This study highlights the importance of MAMs and their role in cellular stress responses, opening the door for therapeutics targeting ER, mitochondria, and MAMs in diseases associated with corneal endothelial degeneration.

## Methods

### Human tissue

Diseased donor corneal endothelium were obtained from Dr. Ula Jurkunas (Massachusetts Eye and Ear, Boston, USA), Dr. Marianne Price, and Dr. Francis Price Jr (Price Vision Group, Indianapolis, USA), and has been used to determine mitochondrial health in their previous publication.^10^ The study was conducted in accordance with the tenets of the Helsinki Declaration of 1975, as revised in 1983, and Institutional Review Board (IRB)/Ethics Committee approval was obtained. Normal donor corneas were purchased from SightLife (Bethlehem, PA). A total of 5 normal donor corneas (65.7 ± 6.7 years) and 5 FECD donor corneas (67.5 ± 8.0 years) were used in the study.

### Cell Culture

Normal immortalized human corneal endothelial cell line (HCEnC-21T) and FECD corneal endothelial cell line (F35T) were generously gifted by Dr. Ula Jurkunas and Dr. Albert Jun (Johns Hopkins University, Baltimore, USA), respectively. 21T and F35T cells were seeded and cultured in Chen’s media as previously described.^11^ Cells were grown on T75 flasks (cat no. 156499; Thermo Fisher) coated with FNC coating mix (cat no. 0407; Athena Enzyme Systems), passaged every two or three days, and maintained in 5% CO_2_ at 37°C. The two cell lines were treated with dimethyl sulfoxide (DMSO) or tunicamycin at varying concentrations, as needed for the experiments.

### MAM Plasmid Transfection

MAMtracker-Green plasmid^18^, used for transfection of 21T and F35T cells, was obtained from Dr. Koji Yamanaka (Nagoya University, Nagoya, Japan). DH5α cells (cat no. C2987H, New England Biolabs) were transformed through heat shock and amplified in media containing kanamycin. Plasmid DNA was isolated using the NucleoSpin Plasmid kit (cat no. 740588, Macherey-Nagel) according to the manufacturer’s protocol. 21T and F35T cells were plated onto 2-well Nunc Lab-Tek II chamber slides (cat. no. 154461, Thermo Fisher) and allowed to reach 70% confluency. Cells were transfected with 0.5 µg plasmid DNA, delivered via Lipofectamine RNAiMAX (cat no. 13778, Invitrogen), for 30 hours, after which treatments (DMSO or tunicamycin) were applied. Transfected and treated cells were fixed in 4% paraformaldehyde (PFA), counterstained with DAPI (cat no. D1306, Invitrogen), and visualized on a confocal microscope (Leica STED). To quantify fluorescence intensity of the MAM plasmid, 15-20 cells were randomly selected per condition, with each condition having 3-6 replicates. MAMtracker-Green plasmid fluorescence intensity was quantified using ImageJ software, corrected for background signal, and normalized against cell area.

### MAM Isolation

21T and F35T cells were plated and expanded on 150 mm culture dishes (cat no. 430599, Corning) for subcellular fractionation. For a single round of MAM isolation, thirty 150 mm cell culture plates containing a total of 600 × 10^6^ cells were used. For baseline conditions, both cell lines were left untreated and allowed to reach 100% confluency before isolation. Treated cells were subjected to either tunicamycin (TUN 1 µg/mL) or dimethyl sulfoxide (DMSO 0.02%) treatment for 24 hours when 70% confluency was reached. Post-treatment, cell pellets for each condition were obtained and processed via the MAM isolation protocol.^19^ A pictorial description of the subcellular fractionation process is shown in Figure 2A, and all centrifugation steps with speeds exceeding *12,000 x g* were performed in a Beckman Coulter Ultracentrifuge (Beckman Ti70.1 rotor). Cell samples from the thirty confluent 150 mm plates were split into 6 equal portions to maximize isolation efficiency. The samples were briefly resuspended in Vance isolation buffer (250 mmol/L mannitol, 5 mmol/L HEPES, pH 7.4, and 0.5 mmol/L EGTA) and gently homogenized in a Dounce homogenizer with 5 strokes using a loose Potter-Elvehjem grinder and 15 strokes using a tight Potter-Elvehjem grinder (Kontes). The homogenate was centrifuged for 10 minutes at *1000 x g*, yielding the nuclear fraction and the supernatant that was collected. The nuclear fraction was reprocessed via resuspension in Vance buffer, and centrifuged again for 10 minutes at *1000 x g*. The resulting pellet can be discarded while the supernatants obtained from both spin steps were pooled and further centrifuged for 15 minutes at *12,000 x g* to obtain crude mitochondrial pellets. The resulting new supernatant was further centrifuged for 60 minutes at *100,000 x g* to separate ER and cytosolic fractions. The crude mitochondria pellets were pooled, then resuspended in a small volume of Vance buffer and loaded onto a 30% Percoll gradient for centrifugation at *95,000 x g* for 30 minutes. This centrifugation step yields separate protein layers in the Percoll gradient, where the upper layer was carefully removed. This upper layer was centrifuged for 10 minutes at *12,000 x g* to obtain the MER (a combination of ER and mitochondrial fractions) pellet and a supernatant. The remaining supernatant was spun at *100,000 x g* for 60 minutes, yielding the MAM fraction and a supernatant that was discarded. The lower protein layer from the Percoll gradient was centrifuged for 10 minutes at *12,000 x g* to collect the free mitochondria fraction and a supernatant (discarded). Samples of all subcellular fractions were collected and stored for further analysis.

### Western Blot

Protein concentrations of isolated subcellular fractions were determined using the Pierce BCA Protein Assay Kit (cat no. 23227, Thermo Fisher). Subcellular fractions were processed with an appropriate volume of LDS Sample buffer and a reducing agent, then separated on 4-12% SDS-PAGE gels and transferred to PVDF membranes. Immunoblotting procedure was performed as previously described.^11^ The following antibodies were used for immunoblotting: PDZD8 (cat no. PA5-46771, Invitrogen), Ero1-Lα (cat no. 3264, Cell Signaling Technology), VDAC (cat no. 4661, Cell Signaling Technology), cytochrome C (cat no. 4272, Cell Signaling Technology), PERK (cat no. 3192, Cell Signaling Technology), Parkin (cat no. 4211, Cell Signaling Technology), GAPDH (cat no. 97166, Cell Signaling Technology), actin (cat no. 3700, Cell Signaling Technology), HRP-linked anti-mouse (cat no. 7076, Cell Signaling Technology) and anti-rabbit (cat no. 7074, Cell Signaling Technology) secondary antibodies.

### Immunocytochemistry

21T and F35T cells were plated onto 2-well chamber slides (cat no. 80286, ibidi), after which appropriate treatments were applied when slides reached 70% confluency. Using our previously published methods^11^ for mitochondrial staining, treated cells were incubated with MitoTracker Deep Red FM (cat no. M22426, Invitrogen) for 30 minutes prior to fixation in 4% PFA. Fixed cells were incubated overnight with primary antibodies, washed, stained with secondary antibodies the following day, and then counterstained with DAPI before storing in PBS at 4°C, as previously published.^11^ Antibodies used in this application include calnexin (cat no. 2679, Cell Signaling Technology), PTDSS1 (cat no. PA5-112828, Invitrogen), and goat anti-rabbit Alexa Fluor 488 (cat no. ab150077, Abcam). All prepared slides were visualized on a confocal microscope (Leica STED). To quantify the fluorescence intensity of PTDSS1 protein expression, 10-20 cells were selected at random per image for each tested condition, with 3-5 images collected per condition. Fluorescence intensity of PTDSS1 expression was quantified using the built-in Analyze/Measure feature in ImageJ, corrected for background signal, and normalized against cell area.

### Transmission Electron Microscopy

Healthy and diseased donor corneal endothelium were processed for transmission electron microscopy (TEM) as published.^11^ Briefly, specimens were rinsed in cacodylate buffer, post-fixed in 2% osmium tetroxide, and then processed for cutting ultra-thin sections (8 μm). Sections were imaged using an FEI Tecnai G2 Spirit transmission electron microscope (FEI, Hillsboro, Oregon) at 80 kV interfaced with an AMT XR41 digital CCD camera (Advanced Microscopy Techniques, Woburn, MA). For the quantification of the average ER-mitochondria contact distance, 3-5 images of control or Fuchs’ tissues were considered. For each tissue condition, 15-20 ER-Mitochondria contact distances were measured and averaged by a blinded investigator using ImageJ software, with a maximum distance of 150 nm, as previously reported in other tissues.^20^ To quantify the percentage of mitochondria-ER interactions, the same images described earlier in this section were used. ER-mitochondria interactions were considered present when contact distance between the two organelles was ≤ 30 nm and were exclusively measured in regions where whole organelles were shown.

### Statistical Analysis

Statistical analysis for comparisons between two groups were assessed using Welch’s and unpaired Student *t*-test on GraphPad Prism. All immunoblot data representations are plotted as mean ± standard error of the mean (SEM) and significance values were set at *\*p*<*0.05, **p*<*0.01, ***p*<*0.001, ****p*<*0.0001*.

## Results

### ER-Mitochondria contact sites, or MAMs, are altered in Fuchs’ corneal endothelial tissues

Mitochondria and ER contacts (also called ER-mitochondria contact sites or MAMs) were first visualized via TEM in normal and Fuchs’ human corneal endothelial tissues to investigate possible structural changes within the MAM with disease progression. A high density of normal mitochondria and ER was observed in healthy control corneal endothelium (Figure 1A), while swollen ER and mitochondrial organelles were detected in the diseased Fuchs’ sample, findings in line with observations previously reported.^10^ Notedly, multiple autophagic/mitophagic vacuoles were also seen in the Fuchs’ tissues (Figure 1A). From past studies, we know that average distance between the ER and mitochondria in a normal cell is 20-30 nm. When visualizing MAMs, the distance between the two organelles, as indicated by red dashes in Figure 1A, was significantly increased to approximately 80 nm in the diseased tissue (Figure 1B). Further analysis of normal and Fuchs’ human corneal endothelium indicated, on average, a 50% reduction in interaction between the ER and mitochondria, suggesting an overall altered MAM organization and interaction in the diseased state (Figure 1C).

**Figure 1.**
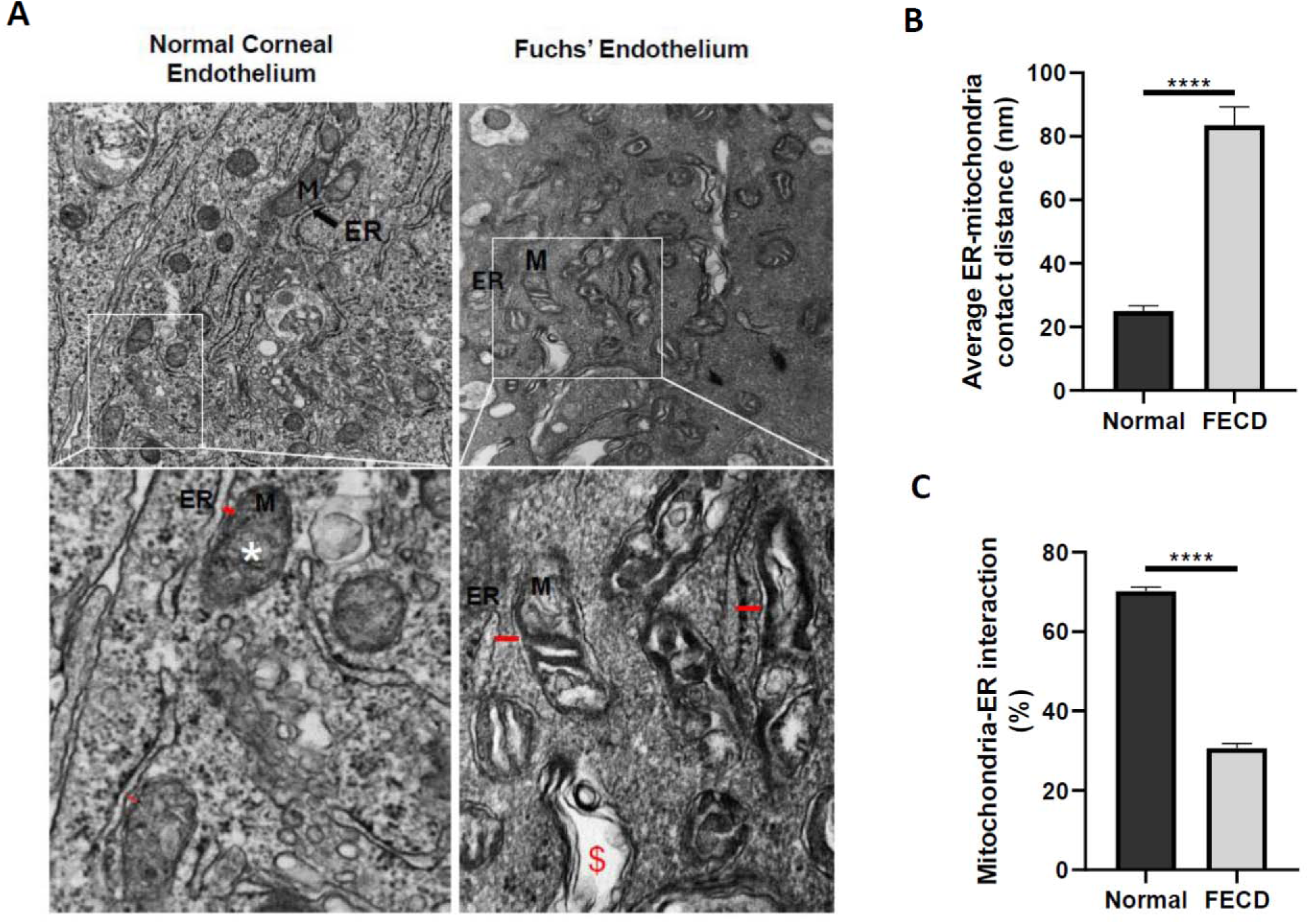
Disruptions of ER-Mitochondria interactions/MAMs in Fuchs’ compared to normal human corneal endothelial tissues. **(A)** TEM images showing swollen morphology of ER and mitochondria with increased distances between the two organelles (marked by red dashes) in Fuchs’ compared to normal human corneal endothelial tissue. Mitophagic vacuoles containing damaged mitochondria are represented as $ (scale bar: 1 μm). **(B)** Bar graph showing increased distance between ER and mitochondria in Fuchs’ compared to normal control tissues (n = 3-5, ****p < 0.0001, unpaired t-test). **(C)** Bar graph also demonstrating decreased percentage of mitochondria-ER interactions in Fuchs’ compared to normal control tissues (n = 3-5, ****p < 0.0001, unpaired t-test).

### ER and mitochondria interactions are disrupted in response to ER stress induction

As significant changes to the MAM domain were observed between normal and Fuchs’ tissues, we next investigated the possible causes of this MAM disruption. Our previous study on ER-mitochondria crosstalk in corneal endothelial cells demonstrated chronic ER stress disrupting mitochondrial functions, thereby contributing to CEnC apoptosis. ^11^ To explore whether this altered ER-mitochondria crosstalk affects MAMs under chronic ER stress conditions, we once again visualized ER-mitochondria contact sites through transmission electron microscopy in 21T cells treated with tunicamycin, a chemical ER stressor blocking proper glycosylation of proteins, at 10 µg/mL for 24 hours. Trends seen in TEM analysis between healthy and diseased human corneal endothelium were similarly observed in HCEnCs subjected to tunicamycin (Figure 2A). Heavily swollen ER and mitochondria alongside enlarged distances (marked by red dashes) between the two organelles were visually detected (Figure 2A). Subsequent analysis of contact distance and organelle interaction between mitochondria and ER revealed the same increased distance and reduced inter-organelle interaction in the ER stressed condition compared to control (Figure 2B, 2C). This reduction in MAMs in response to ongoing ER stress was further confirmed in a qualitative colocalization study of 21T HCEnCs subjected to tunicamycin at 10 µg/mL for 24 hours, labelling ER and mitochondria with calreticulin and MitoTracker, respectively (Figure 2D). Visualized through confocal microscopy, a substantial decrease in overlap of ER and mitochondrial labelling probes were detected through color shift (less yellow – reduced overlap) in ER-stressed 21T cells as opposed to the more widespread colocalization of ER and mitochondrial probes (more yellow – more overlap) in the vehicle control condition (Figure 2D). This finding, in addition to the quantitative data of the TEM analysis between control and tunicamycin-treated 21T cells, suggests interactions between ER and mitochondria, as well as their contact sites, diminish in response to chronic ER stress in corneal endothelial cells.

**Figure 2.**
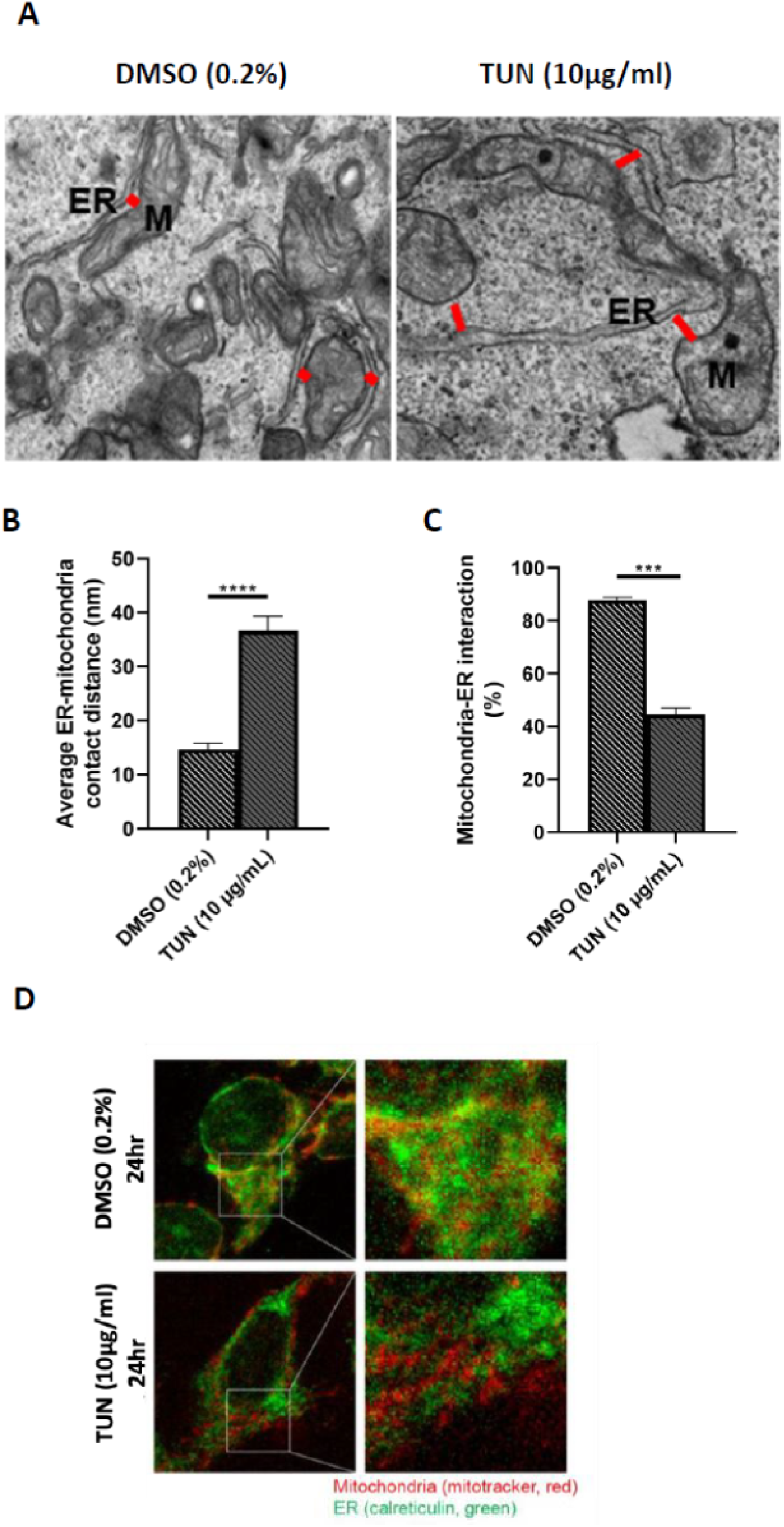
ER-Mitochondria contact sites/MAMs are altered in response to ER stress induction. **(A)** TEM showing significant swelling of ER and mitochondria (M) as well as an increased distance between these two organelles in HCEnC-21T after treatment with tunicamycin (10 mg/ml) for 24 hours compared to control (DMSO) (scale bar: 0.5 μm). **(B)** Bar graph demonstrating enlarged ER-mitochondria contact distance and **(C)** reduced percentage of mitochondria-ER interactions in HCEnC-21T under tunicamycin treatment (10 mg/ml, 24 hours) compared to DMSO control (n = 3-5, ***p < 0.001, ****p < 0.0001, unpaired t-test). **(D)** Immunostained images of HCEnC-21T labelled with mitochondria marker (MitoTracker) in red and ER marker (calreticulin) in green show a decrease in co-localization of mitochondria and ER, represented as yellow, in 21T after treatment with tunicamycin (10 μg/ml) for 24 hours compared to vehicle control.

### Disruptions to MAMs in response to chronic ER stress are time and dose-dependent in corneal endothelial cells

Having detected alterations to ER-mitochondrial contacts post-ER stress exposure, we sought to further explore how chronic ER stress affects MAM distance and interaction. The association between mitochondria and ER was investigated using a transfected MAMtracker-Green plasmid that encodes for dimerization-dependent green fluorescent proteins (GFP) conjugated to components localizing separately to the ER and mitochondria, only emitting fluorescence when heterodimers (ER-localized and mitochondria-localized) coalesce.^18^ With our previous study on ER-mitochondria crosstalk showcasing differences in cellular response across time^11^, we applied tunicamycin treatment to 21T cells and tracked temporal changes in MAM interactions using the MAMtracker-Green plasmid. Plasmid-transfected 21T cells exposed to 1 µg/mL of tunicamycin was visualized under confocal microscopy at 6 hours, 24 hours, and 48 hours of stressor treatment (Figure 3A, 3B, 3C). Mean fluorescence intensity analysis across the three time points revealed a significant elevation in MAMtracker-Green plasmid fluorescence intensity at 6 hours of stressor exposure compared to DMSO (Figure 3D), followed by a plateau in intensity difference, where no statistical significance was detected between the stressed and control conditions at 24 hours of tunicamycin treatment (Figure 3E). This leveling out in fluorescence intensity of the tracker plasmid was concluded by a sharp decline in detected fluorescent signal in the ER-stressed condition of 21T cells after 48 hours of exposure (Figure 3F), suggesting that over time, ER-mitochondria contacts experience an increase in connection in response to acute ER stress, then gradually distance and lose association as the stressor is continued.

**Figure 3.**
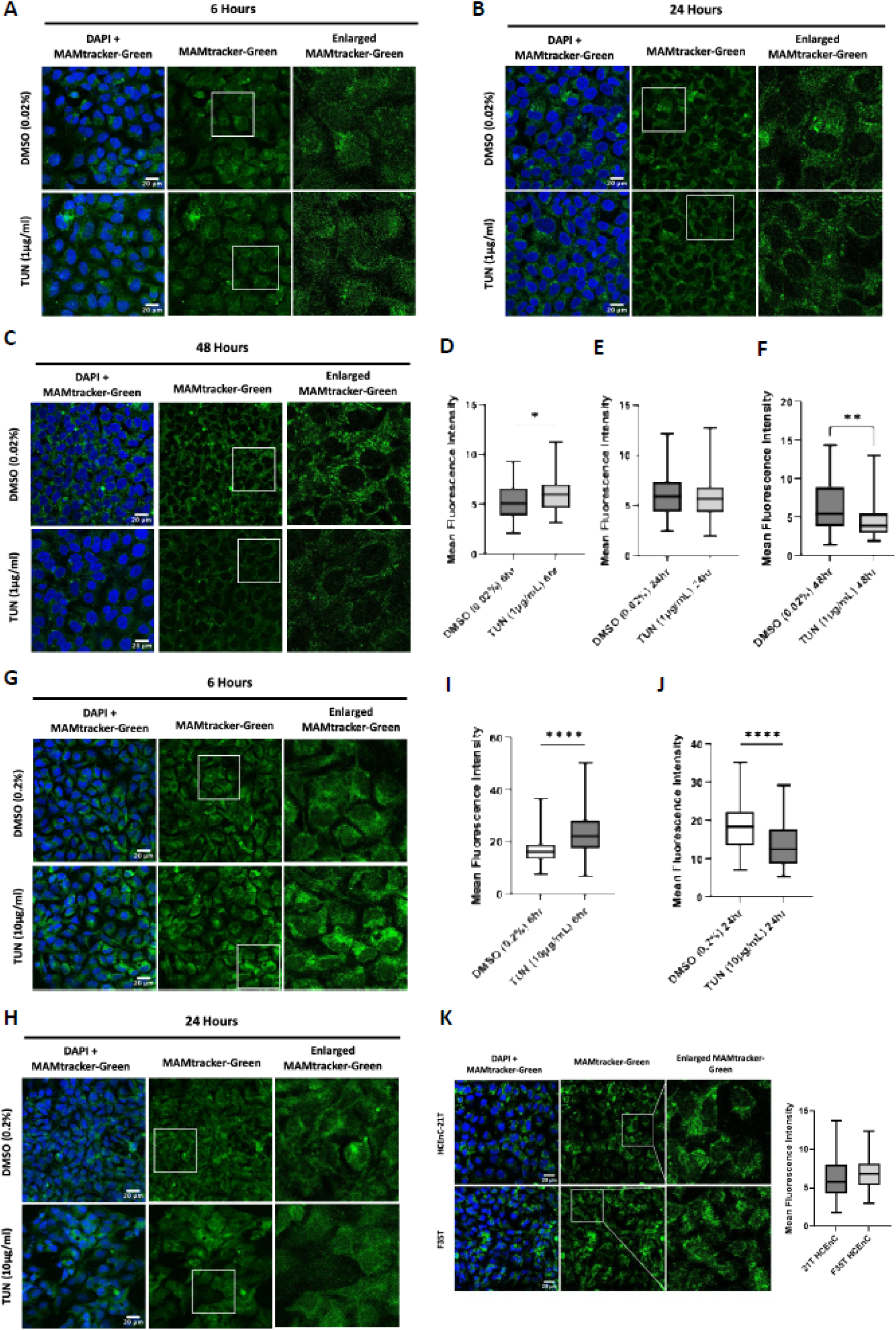
Disruption of ER-mitochondria interaction in response to ER stress is time- and dose-dependent. **(A-C)** Immunostained images of MAM tracker-Green (green) transfected 21T human corneal endothelial cells, nuclei counterstained with DAPI (blue), treated with tunicamycin 1 µg/mL for 6, 24, and 48 hours showed increased mitochondria-ER contact as early as 6 hours, subsequently equalizing at 24 hours, then decreased contact at 48 hours of ER stress treatment (scale bar: 20 µm). **(D-F)** Quantification of fluorescence intensity show a significant elevation of plasmid fluorescence at 6 hours post-tunicamycin treatment compared to control, while no statistical significance was found between the stressed and control conditions at 24 hours. A drop in fluorescence intensity was observed at 48 hours post-tunicamycin exposure (n=4-6, *p < 0.05, **p < 0.01, Welch’s t-test). **(G-H)** Representative immunostained images of MAMtracker-Green (green) transfected 21T cells, nuclei counterstained with DAPI (blue) subjected to a higher dose of tunicamycin at 10 µg/mL for 6 and 24 hours. A similar trend of increased MAM contact at 6 hours was seen while a sharp decline in interaction was observed at 24 hours (scale bar: 20 µm). **(I-J)** Box plots of mean fluorescence intensity quantifications revealed a greater rise in intensity signals at 6 hours post-ER stress when compared to control. A significant decrease in fluorescence intensity was detected at 24 hours of tunicamycin treatment compared to that seen in the control condition at the same time point (n=6, ****p < 0.0001, Welch’s t-test). **(K)** Immunostained images of MAM tracker-Green (green) transfected 21T and F35T cells, nuclei counterstained with DAPI (blue), showing no significant difference in intensity, suggesting similar levels of ER-mitochondria interactions/ MAMs in both cell lines at baseline (non-stressed) (scale bar: 20 µm, n=4).

Increasing the dose of stressor applied yielded a familiar response in MAMtracker-Green plasmid fluorescence intensity. Treatment of 21T HCEnCs with 10 µg/mL of tunicamycin also demonstrated an initial upsurge of detected fluorescent intensity signal in the stressed condition (Figure 3G,3I), however, by the 24 hours mark, rather than stabilizing like the lower dose counterpart, a significant reduction of fluorescent signal was found between the tunicamycin-treated cells and the vehicle control cells (Figure 3H, 3J). This slight alteration in trend appears to indicate that in addition to temporal differences, MAM interactions also respond to dose differences in the applied stressor. Compared to the MAMtracker-Green plasmid fluorescence intensity of its lower dose counterparts (1 µg/mL), the fold changes in mean fluorescence intensity between stressed and control conditions at both 6 and 24 hours of the 10 µg/mL tunicamycin dose are more drastic; at 6 hours of tunicamycin treatment, fold change in intensity value between vehicle control and ER-stressed conditions are 1.15 and 1.37 for the 1 µg/mL dose and 10 µg/mL dose, respectively, while at 24 hours of stressor exposure, fold change in fluorescence intensity between DMSO and tunicamycin conditions are 0.95 and 0.71 for the 1 µg/mL dose and 10 µg/mL dose, respectively. These alterations in ER and mitochondrial interactions in response to strength and duration of stressor exposure reflect a time-and dose-dependency amongst the contact sites between the two organelles, further highlighting the dynamic nature of this MAM domain and the ongoing shifts that occur within its interactions.

Interestingly, further exploration of MAM contacts using the MAMtracker-Green plasmid revealed a surprising result in the interaction between ER and mitochondria in 21T and F35T cells at baseline. When probing for tracker plasmid fluorescence signals, no significant differences were found in MAM interactions amongst the two cell lines when left untreated (Figure 3K).

### Chronic ER stress alters expression of MAM functional protein in corneal endothelial cells

Given the observed dynamic changes in ER-mitochondria contact sites, we further investigated the effects of chronic ER stress on more functional aspects of the MAM. MAM is heavily involved in several metabolic processes necessary for cellular homeostasis, including facilitating lipid transport between the mitochondria and ER, particularly in the production and movement of phospholipids.^22,23^ One way to determine MAM function is through synthesis and transport of phospholipids^24^ Altered MAM structure often leads to decreased MAM function, as measured by the transport of MAM phospholipid enzymes, including phosphatidylserine synthase (PTDSS1) and phosphatidylserine decarboxylase (PISD). Briefly, phosphatidylserine is synthesized in the MAM domain by PTDSS1, and is then translocated to the mitochondria, where phosphatidylserine is decarboxylated by PISD.^24^ PTDSS1 is a protein highly enriched in the MAM fractions. ^25,26–28^ We first focused on detecting PTDSS1 expression in 21T and F35T cells under basal conditions using immunofluorescence and found elevated expression of PTDSS1 in F35T cells compared to 21T HCEnCs, as seen through a significantly higher mean fluorescence intensity value (Figure 4A, 4B). While no difference in ER-mitochondria contact was uncovered between 21T and F35T cells at baseline when probing with the MAMtracker-Green plasmid, there appears to be a variation in functional protein expression present at the MAM domain between the two cell lines. Continuing in our exploration of possible effects continuous ER stress has on MAM functions, PTDSS1 expression was investigated in 21T cells subjected to ER stress (tunicamycin 1 μg/mL) for 6 hours (Figure 4C). This initial timepoint did not reveal vast changes in phosphatidylserine synthase expression amongst the tested conditions (Figure 4D), suggesting at acute levels of ER stress, the degree of functional PTDSS1 proteins present at the MAM are not significantly altered. As ER-mitochondria interactions are evident to change with time of stress exposure, it was important to observe protein expression of PTDSS1 when ER stress was prolonged. 21T HCEnCs subjected to tunicamycin treatment at 1 μg/mL for 24 hours revealed a drop in PTDSS1 antibody fluorescence intensity compared to cells subjected to DMSO vehicle control (Figure 4E, 4F). The decrease in detected fluorescence intensity value of the ER-stressed condition indicates lower expression of PTDSS1 protein at various ER-mitochondria contact sites, potentially signifying a decrease in MAM function with extended ER stressor treatments, correlating with the decline in MAM contacts seen at later time points of tunicamycin treatment (Figure 3C, 3H).

**Figure 4.**
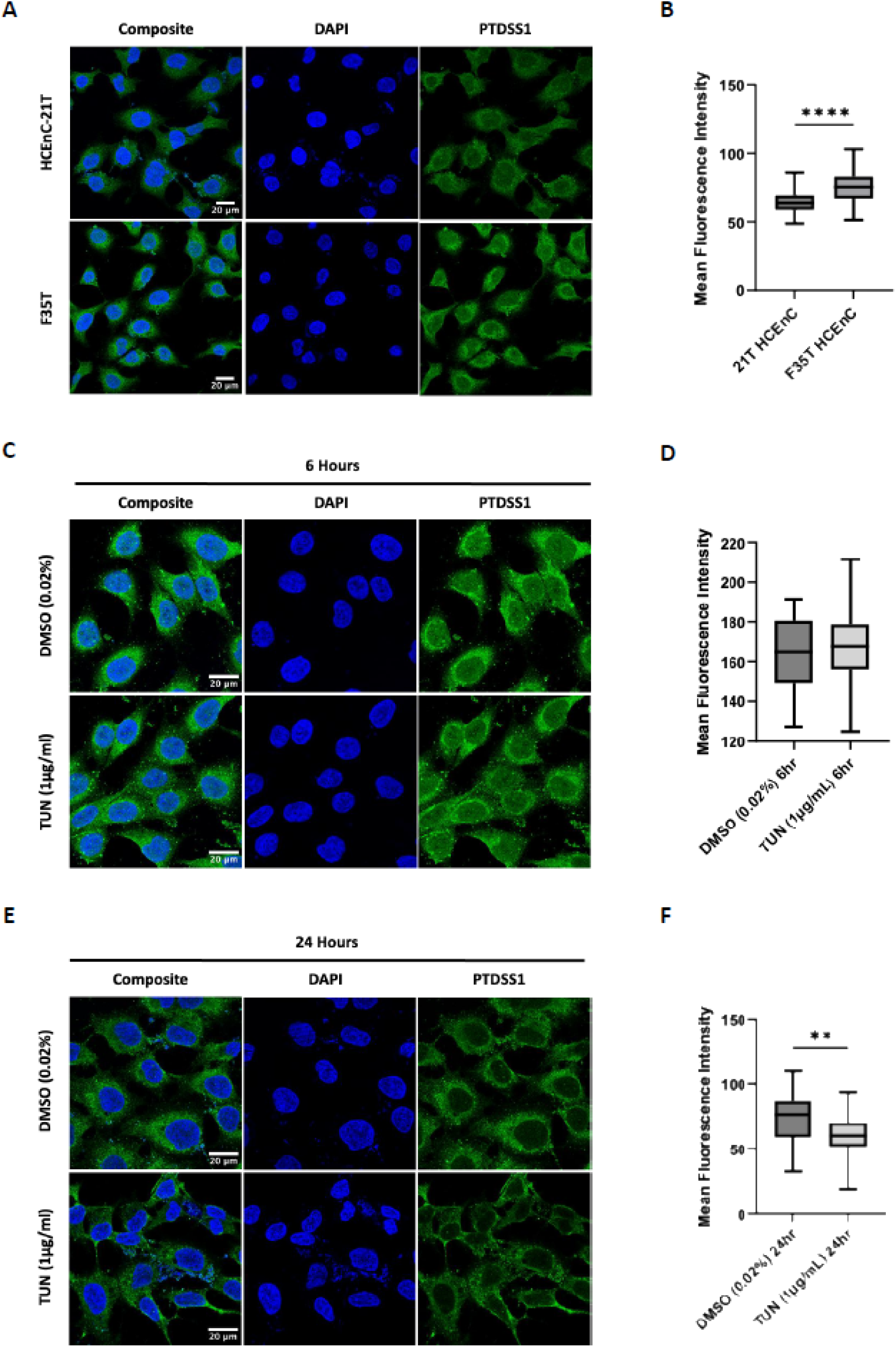
Continuous ER stress alters expression of MAM functional protein in human corneal endothelial cells. **(A)** Representative immunostained images of MAM protein PTDSS1 (phosphatidylserine synthase enzyme; green) in untreated 21T and F35T corneal endothelial cells, nuclei counterstained with DAPI (blue) (scale bar: 20 µm). **(B)** Box plot demonstrating increased mean fluorescence intensity of PTDSS1 in diseased F35T cell line compared to healthy 21T HCEnCs at baseline (n=8-9, ****p < 0.0001, Welch’s t-test). **(C-D)** Representative immunostained images of PTDSS1 (green) in 21T cells, nuclei counterstained in blue with DAPI, following exposure to ER stressor tunicamycin for 6 hours (scale bar: 20 µm). Quantification showed no difference in fluorescence intensity between stressed and control conditions at this time point (n=4). **(E)** Representative immunostained images of 21T cells showing reduced PTDSS1 fluorescence signal (green), nuclei counterstained with DAPI (blue), after treatment with tunicamycin 1 µg/mL at 24 hours compared to DMSO control (0.2%). **(F)** Box plot demonstrating decreased fluorescence intensity of PTDSS1 for tunicamycin-treated 21T cells compared to vehicle control (n = 4, **p < 0.01, Welch’s t-test).

### Stress response proteins, PERK and Parkin, accumulate in the MAM domain post-ER stress induction

To better under molecular and functional changes that occur between the ER and mitochondria and their contact sites, we isolated MAMs, mitochondria, and ER organelles from 21T and F35T cell lines. Figure 5A illustrates the schematic steps involved in isolating the various fractions, including MAMs. For subcellular fractionation, we isolated total homogenate after centrifugation of whole cell lysate at *1,000 x g* and the mitochondrial fraction was collected post-centrifugation at *12,000 x g*. Further separation of MAM and ER fractions were obtained after ultracentrifugation at speeds indicated in Figure 5A, through the use of a Percoll gradient. The isolated subcellular fractions were verified by demonstrating enrichment of fraction-specific proteins (Figure 5B, 5C) as previously published.^21^ PDZD8, a recognized marker of the MAM domain, was observed to be enriched in the MAM fraction of both 21T and F35T cells (Figure 5B, 5C). The MAM protein marker was also seen in the crude mitochondria fraction of both cells lines as crude mitochondria consists of a mixture of pure mitochondria organelles and the mitochondria-associated membranes (MAMs) (Figure 5B, 5C). Ero1-Lα, a luminal ER protein, has the highest subcellular concentration in ER fractions of both cell lines, with trace degrees of localization within mitochondria and MAM fractions (Figure 5B, 5C). VDAC and cytochrome C, both mitochondrial proteins, were mainly found within mitochondrial fractions of 21T and F35T cells, with faint traces of VDAC seen in the MAM fraction being that it also acts as tether proteins within complexes inside the MAM domain (Figure 5B, 5C).^39^

**Figure 5.**
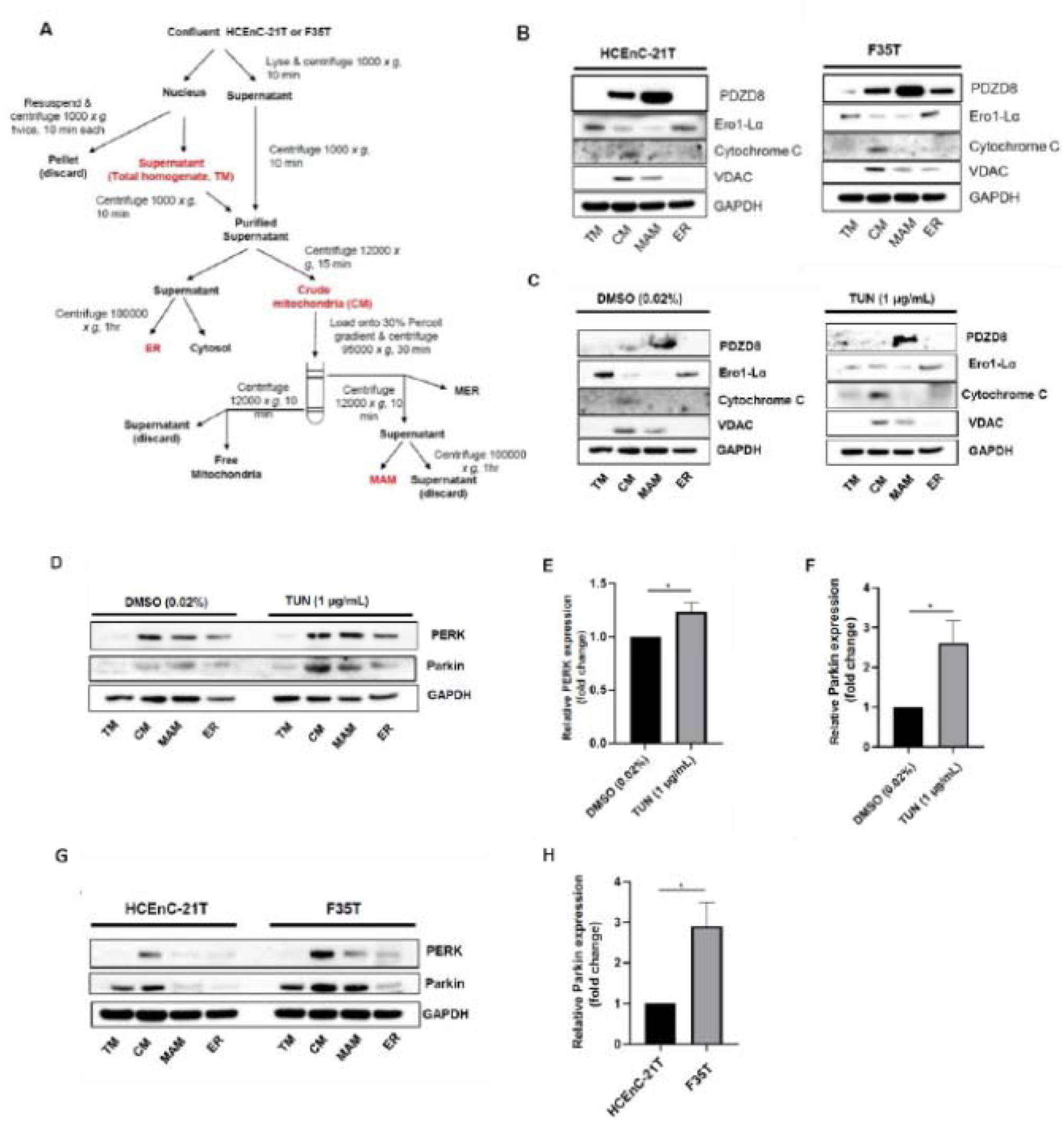
Accumulation of ER stress response protein, PERK, and mitochondria damage sensor protein, Parkin, in the MAM domain after continued ER stress. **(A)** Description of subcellular fractionation method for mitochondria-associated ER membrane (MAMs), endoplasmic reticulum (ER), and crude mitochondria (CM) isolation in corneal endothelial cells. **(B)** Representative Western blot data confirming subcellular fraction markers PDZD8 (MAM marker), Ero1-Lα (ER marker), VDAC and cytochrome C (mitochondrial markers), and GAPDH (loading control) in normal 21T and diseased F35T cells. **(C)** Representative Western blots confirming subcellular fraction markers mentioned previously in 21T cells after treatment with control DMSO (0.02%) or ER stressor tunicamycin (1 µg/mL) at 24 hours. **(D)** Representative Western blots showing increased expression of PERK and Parkin proteins in the MAM fraction in 21T cells after exposure to ER stressor tunicamycin (1 µg/mL) for 24 hours compared to control DMSO (0.02%). **(E-F)** Bar graphs demonstrating an increase in fold expression of PERK and Parkin proteins under stressed condition compared to control (n = 4, *p < 0.05, unpaired t-test). **(G)** Representative Western blots of subcellular fractions in 21T and F35T cell line at baseline (untreated), probing for PERK, Parkin and GAPDH. **(H)** Bar graph demonstrating increased Parkin expression in the MAM fraction of diseased F35T corneal endothelial cells compared to healthy 21T cells at baseline (n = 3, *p < 0.05, unpaired t-test).

After a CEnC MAM fraction was established and confirmed, a comprehensive list of stress response proteins was probed and within the group, PERK and Parkin where the two proteins exhibiting the most significant alterations within the MAM fraction with continued ER stress induction (Figure 5D, 5G). The expression level of PERK, an ER stress sensor protein, was significantly elevated in the MAM fraction of 21T cells treated with tunicamycin at 1 μg/mL for 24 hours (Figure 5D, 5E). An accumulation of Parkin, a ubiquitin ligase sensing mitochondrial damage, was similarly seen in 21T MAM fraction following 24 hours of ER stress exposure (1 μg/mL) (Figure 5D, 5F). The finding of both ER and mitochondria stress proteins within this MAM domain after stress induction suggests a possible region of interaction between ER and mitochondria stress signaling, linking the two organelles and allowing for crosstalk to occur. Probing for PERK and Parkin within the MAM fractions of 21T and F35T cell lines at baseline revealed upregulation of Parkin protein expression in the MAM domain of the diseased F35T cell line (Figure 5G, 5H). While no statistical significance has yet been detected for elevated PERK expression in F35T cells compared to healthy 21T HCEnCs, an increasing trend of increased protein expression can be seen for PERK in the MAM of F35T cells (Figure 5G), suggesting that ER and mitochondrial stress-sensor proteins, such as PERK and Parkin, accumulate in the MAM and may play a role in mediating cellular responses to continuous ER stress.

## Discussion

Over the last decades, ER^6,29,30^ and mitochondria stress^31–33^ have been widely studied and implicated in CEnC apoptosis in FECD. Our group first identified crosstalk between ER and mitochondria in the corneal endothelium, previously reporting on chronic ER stress contributing to mitochondrial bioenergetic and dynamic alterations, ultimately leading to the activation of apoptotic pathways in CEnCs.^11,34,35^ However, there remains a need to explore the location of this interaction, and this study attempts to shed light on the site of ER-mitochondrial crosstalk, specifically, at ER-associated mitochondria membranes (MAMs). A recent study had begun to describe altered mitochondria-associated membranes in Fuchs’, highlighting an increase in ER-mitochondria interactions in the diseased state.^36^ While this new study touches upon a few important attributes of MAMs in FECD, using whole cell lysates to probe for MAM proteins and colocalization studies of ER and mitochondria proteins to visualize interaction ^36^, the study of mitochondria-associated membranes in corneal endothelial cells can benefit from more direct and refined analysis into the core of the subject: the MAM domain itself. This missing information is the aim our study attempts to fill, employing advanced techniques available, including subcellular fractionation to isolate MAMs,^37,38^ electron microscopy,^39,40^ and MAM tracker plasmids,^24^ as performed previously in other studies, to describe MAM changes in Fuchs’. Another important distinction to make is our study’s objective to provide a temporal overview of the alterations to ER-mitochondrial interactions in the MAM of Fuchs’ corneal endothelium, a vital element to consider due to the degenerative nature of the disease and the major contributions of sustained/chronic stress on the progression of FECD.^1,6,9^

TEM imaging analysis of human tissues in this study demonstrated an increase in ER-mitochondria contact distance, as well as decreased associations of ER and mitochondria, in human Fuchs’ tissues compared to the healthy control, suggesting a reduction in MAM contacts in the diseased condition. However, this contrasts with reports of increased MAMs found in human Fuchs’ tissues^36^ and in Alzheimer’s disease.^41^ This discrepancy could result from different disease stages of the patients at the time of tissue collection, as both Fuchs’ and Alzheimer’s are degenerative diseases with variations in severity.^1,41^ The diseased human samples used in this study presented with numerous autophagic/mitophagic vacuoles, and in combination with the observation of swollen mitochondria in the Fuchs’ tissues, suggests that the patient had reached the stage where mitophagy had been induced. The collection of late-stage Fuchs’ tissues presents a limitation to our study, and in the future, we will investigate structural changes in MAMs using human tissues obtained at various stages of Fuchs’ progression.

Between normal (21T) and diseased (F35T) cell lines, no significant difference in MAMtracker-Green fluorescence intensity was observed, indicating no difference in ER-mitochondria contacts at the basal level. Though initially surprising, considering the changes witnessed between normal and Fuchs’ corneal endothelial tissues, further analysis appears to indicate this lack of change not as unexpected. Our group has previously investigated basal differences between 21T and F35T cell lines in regards to ER and mitochondrial integrity, and had found a similar lack of change in mitochondrial membrane potential despite seeing an elevation in apoptotic and ER stress markers in the diseased cell line.^11,35^ A significantly identifiable difference, however, was a substantial drop in mitochondrial energy generation, where the rate of ATP production in F35T cell line was far lower than that of the healthy 21T cell line, possibly contributing to the elevation of apoptotic markers detected in F35T cell line.^35^ The information provided by this previous investigation suggests the presence of functional disruptions rather than structural differences between F35T and 21T cells. When taking into consideration the revelation of discrepancies in expression of MAM functional protein, PTDSS1, between 21T and F35T HCEnCs (Figure 4A, 4B), it appears that functional changes are more prominent and sensitive to detection in immortalized F35T cells, rather than structural alterations. While, this may be one possible explanation of the differences exhibited between the two cell lines at baseline, it is also important to note that the cell lines have been immortalized and may exhibit differences from their primary forms, representing another limition to this study. Future investigations would benefit from the use of primary healthy and FECD HCEnCs to visualize ER and mitochondria interaction modifications for better representation of intracellular changes.

Exploration of the impact of ER stress on MAMs showed an increase in contact between the ER and mitochondria, as evidenced by increased MAMtracker-Green fluorescence intensity within 6 hours of ER stress treatment (Figure 3A, 3G). This finding is in line with reports from a recent MAM study, highlighting a similar increase in contact in immortalized human corneal endothelial cell line after ER stress exposure.^36^ Our findings of an initial increase in ER-mitochondria contact are further confirmed by another study in HeLa cells ^42^, where increased colocalization of the two organelles was observed in response to the early onset of chronic ER stress. Increased MAMs during the beginning hours of ER stress exposure are considered an early compensatory response to the elevated metabolic demands associated with ER stress response.^42^ Further study into the effects of chronic ER stress on MAMs demonstrated a leveling out of ER-mitochondria contact at 24 hours of stress exposure (1 µg/mL), as seen through the lack of significant difference in MAMtracker-Green fluorescence intensity between stressed and control conditions in 21T cells (Figure 3B, 3E). This was followed by a steep drop in plasmid fluorescence at 48 hours of tunicamycin treatment (1 µg/mL) (Figure 3C, 3F), indicating reduced ER-mitochondrial interactions at this later point of ER stress exposure, likely caused by the effects of sustained ER stress over time. When ER stress experienced by CEnCs is prolonged and continuous, the available intracellular stress responses may not be sufficient to overcome the disruptions of the stressor. This then can lead to mitochondrial and ER swelling, as well as fragmentation, ultimately contributing to CEnC apoptosis, as previously published by our lab.^43^ This increased swelling and fragmentation of ER and mitochondria potentially distances the organelles, leading to decreased MAMs under continuous ER stress. Our data on reduced MAMs seen in end-stage Fuchs’ tissues is also in line with the notion that sustained organellar stress results in disrupted MAMs under pathological conditions. Furthermore, our investigation of ER stress and its impacts on MAMs also revealed a dose-dependent element to the MAM response following ER stress induction. Treatment with an elevated tunicamycin dose of 10 µg/mL showed a more rapid progression of ER-mitochondria interaction change over time, where an initial increase in MAM contact was still seen at 6 hours, but an even sharper decline in interaction was evident at 24 hours with a significant decrease in MAMtracker-Green fluorescence intensity seen in the ER stressed condition (Figure 3G-3J). As the dynamic nature of the MAM domain is showcased by the data of this study, with ER and mitochondrial interactions altering based on stressor dosing and exposure time, past studies of ER stress and MAMs in FECD could be further elaborated with explorations into this phenomenon, as past reports of increased MAMs following ER stress in HCEnCs are based only on high dose tunicamycin treatments (30 µg/mL) at acute times of exposure (9 hours).^36^

To further understand the role of chronic ER stress in MAM organization, we isolated subcellular fractions from 21T cells post-tunicamycin treatment and observed both PERK and Parkin expressions to be significantly upregulated in the MAM fraction after 24 hours of stress exposure (Figure 5D-5F). A similar significant upregulation of Parkin, and an upward trend in PERK expression, was also seen in the MAM fraction of F35T cells compared to 21T HCEnCs (Figure 5G, 5H), suggesting possible involvement of these stress response proteins in the regulation of MAMs and in the communication of stress signals between the ER and mitochondria within FECD. In line with our data, PERK enrichment in the MAM domain has been previously reported in murine embryonic fibroblasts, and had played a role in relaying reactive oxygen species (ROS) from the ER to mitochondria, contributing to apoptosis during chronic ER stress.^14^ Similarly, Parkin protein has been previously shown to mediate mitophagic processes that help regulate cardiac MAMs during endotoxemia ^44^, while human fibroblasts deficient in Parkin was shown to exhibit decreased MAMs.^17^ Conversly in HeLa cells, enhanced Parkin expression was found to promote ER-mitochondria crosstalk via Ca^2+^ transfer, again suggestive of Parkin’s role in maintaining ER-mitochondria interaction.^45^ Though our present indicates a possible involvement of PERK and Parkin, the mechanism by which the stress response proteins regulate MAMs under continuous stress remains unclear and is another limitation of this study, which will be investigated in the future to elucidate the mode of MAM disruption by Parkin and PERK under ER stress.

MAMs regulate various physiological functions, including calcium transport and lipid exchange, thereby making the disruption of MAMs a contributing factor to the pathogenesis of several metabolic and degenerative diseases. As previously mentioned, a significant upregulation in MAM functional protein PTDSS1 was seen in F35T cells compared to health 21T HCEnCs (Figure 4A, 4B), indicative of potential functional changes in the MAM that are present in the diseased cell line. Interestingly, following treatment of tunicamycin 1 µg/mL at 6 hours revealed no statistically significant alterations to PTDSS1 expression between healthy control and ER stressed 21T cells (Figure 4C, 4D), though this lack of change could possibly be attributed to a delay in protein synthesis response at the initial hours of stress that could potentially catch up at a later time point. Despite this lack of change at 6 hours, treatment of tunicamycin 1 µg/mL at 24 hours displayed a decline in PTDSS1 antibody fluorescence intensity, indicating decreased PTDSS1 protein expression as the ER stress is prolonged (Figure 4E, 4F). With decrease in expression of a functional MAM protein, the data suggests a correlated reduction in MAM function alongside the previously discussed structural disruptions to MAMs in cells subjected to sustained ER stress. A recent study in human osteosarcoma cells lacking mitochondrial DNA also displayed similar alterations in MAM function following decreased enzymatic activity in two proteins that contributed to mitochondrial disease.^24^ While the data is suggestive of a decrease in function of MAM lipid-enzyme activity after sustained ER stress, the influence of other intracellular stressors, such as mitochondrial stress or oxidative stress, in the impairment of MAMs in FECD is still unclear, representing yet another limitation to this study, of which we hope to investigate in the future.

In conclusion, the decrease in MAM contact and function, in combination with the accumulation of ER and mitochondria stress adaptive response proteins in the MAM, suggests that mitochondria-associated ER membranes act as potential hubs for crosstalk between the two organelles in CEnCs, where ER signaling proteins and mitochondrial integrity regulators coalesce to coordinate a global cellular response to stress. The MAM domain remains an area of exciting research, with future studies aimed at better elucidating additional signaling pathways and regulators, as well as explorations into other functional changes that have a larger impact on CEnC viability.

## Author contributions

SL, SK, HOA, SQ, AK, WS, LR, VK performed experiments, analyzed data. SL and VK wrote and revised the manuscript. MP, FP, and UV provided the human tissues and helped in revising the manuscript. HOA, ES helped us standardize MAM experiments and revised the manuscript. VK is responsible for all the funding of the lab.

## Acknowledgments

We sincerely thank Allison Sowa, Bill Janssen for electron microscopy processing and imaging at The Microscopy CoRE, Icahn School of Medicine at Mount Sinai. We also thank Ula V. Jurkunas (Schepens Eye Research Institute, Harvard University) for providing HCEnC-21T cell line and Albert Jun (Wilmer Eye Institute, Johns Hopkins University) for providing F35T cell line.

## Data availability

Data generated in this study will be available from the corresponding author upon request.

## References

1. Iliff, B.W., Riazuddin, S.A., and Gottsch, J.D. (2012). The genetics of Fuchs’ corneal dystrophy. Expert Rev Ophthalmol 7, 363–375. 10.1586/eop.12.39.

2. Krachmer, J.H., Purcell, J.J., Jr., Young, C.W., and Bucher, K.D. (1978). Corneal endothelial dystrophy. A study of 64 families. Arch Ophthalmol 96, 2036–2039. 10.1001/archopht.1978.03910060424004.

3. Zhu, A.Y., Eberhart, C.G., and Jun, A.S. (2014). Fuchs endothelial corneal dystrophy: a neurodegenerative disorder? JAMA Ophthalmol 132, 377–378. 10.1001/jamaophthalmol.2013.7993.

4. Aiello, F., Gallo Afflitto, G., Ceccarelli, F., Cesareo, M., and Nucci, C. (2022). Global Prevalence of Fuchs Endothelial Corneal Dystrophy (FECD) in Adult Population: A Systematic Review and Meta-Analysis. J Ophthalmol 2022, 3091695. 10.1155/2022/3091695.

5. Li, Q.J., Ashraf, M.F., Shen, D.F., Green, W.R., Stark, W.J., Chan, C.C., and O’Brien, T.P. (2001). The role of apoptosis in the pathogenesis of Fuchs endothelial dystrophy of the cornea. Arch Ophthalmol 119, 1597–1604. 10.1001/archopht.119.11.1597.

6. Engler, C., Kelliher, C., Spitze, A.R., Speck, C.L., Eberhart, C.G., and Jun, A.S. (2010). Unfolded protein response in fuchs endothelial corneal dystrophy: a unifying pathogenic pathway? Am J Ophthalmol 149, 194–202 e192. 10.1016/j.ajo.2009.09.009.

7. Okumura, N., Hashimoto, K., Kitahara, M., Okuda, H., Ueda, E., Watanabe, K., Nakahara, M., Sato, T., Kinoshita, S., Tourtas, T., et al. (2017). Activation of TGF-beta signaling induces cell death via the unfolded protein response in Fuchs endothelial corneal dystrophy. Sci Rep 7, 6801. 10.1038/s41598-017-06924-3.

8. Gupta, S., Giricz, Z., Natoni, A., Donnelly, N., Deegan, S., Szegezdi, E., and Samali, A. (2012). NOXA contributes to the sensitivity of PERK-deficient cells to ER stress. FEBS Lett 586, 4023–4030. 10.1016/j.febslet.2012.10.002.

9. Halilovic, A., Schmedt, T., Benischke, A.S., Hamill, C., Chen, Y., Santos, J.H., and Jurkunas, U.V. (2016). Menadione-Induced DNA Damage Leads to Mitochondrial Dysfunction and Fragmentation During Rosette Formation in Fuchs Endothelial Corneal Dystrophy. Antioxid Redox Signal 24, 1072–1083. 10.1089/ars.2015.6532.

10. Benischke, A.S., Vasanth, S., Miyai, T., Katikireddy, K.R., White, T., Chen, Y., Halilovic, A., Price, M., Price, F., Jr., Liton, P.B., and Jurkunas, U.V. (2017). Activation of mitophagy leads to decline in Mfn2 and loss of mitochondrial mass in Fuchs endothelial corneal dystrophy. Sci Rep 7, 6656. 10.1038/s41598-017-06523-2.

11. Qureshi, S., Lee, S., Steidl, W., Ritzer, L., Parise, M., Chaubal, A., and Kumar, V. (2023). Endoplasmic Reticulum Stress Disrupts Mitochondrial Bioenergetics, Dynamics and Causes Corneal Endothelial Cell Apoptosis. Invest Ophthalmol Vis Sci 64, 18. 10.1167/iovs.64.14.18.

12. Markovinovic, A., Greig, J., Martin-Guerrero, S.M., Salam, S., and Paillusson, S. (2022). Endoplasmic reticulum-mitochondria signaling in neurons and neurodegenerative diseases. J Cell Sci 135. 10.1242/jcs.248534.

13. Wang, N., Wang, C., Zhao, H., He, Y., Lan, B., Sun, L., and Gao, Y. (2021). The MAMs Structure and Its Role in Cell Death. Cells 10. 10.3390/cells10030657.

14. Verfaillie, T., Rubio, N., Garg, A.D., Bultynck, G., Rizzuto, R., Decuypere, J.P., Piette, J., Linehan, C., Gupta, S., Samali, A., and Agostinis, P. (2012). PERK is required at the ER-mitochondrial contact sites to convey apoptosis after ROS-based ER stress. Cell Death Differ 19, 1880–1891. 10.1038/cdd.2012.74.

15. Ge, P., Dawson, V.L., and Dawson, T.M. (2020). PINK1 and Parkin mitochondrial quality control: a source of regional vulnerability in Parkinson’s disease. Mol Neurodegener 15, 20. 10.1186/s13024-020-00367-7.

16. McLelland, G.L., Goiran, T., Yi, W., Dorval, G., Chen, C.X., Lauinger, N.D., Krahn, A.I., Valimehr, S., Rakovic, A., Rouiller, I., et al. (2018). Mfn2 ubiquitination by PINK1/parkin gates the p97-dependent release of ER from mitochondria to drive mitophagy. Elife 7. 10.7554/eLife.32866.

17. Basso, V., Marchesan, E., Peggion, C., Chakraborty, J., von Stockum, S., Giacomello, M., Ottolini, D., Debattisti, V., Caicci, F., Tasca, E., et al. (2018). Regulation of ER-mitochondria contacts by Parkin via Mfn2. Pharmacol Res 138, 43–56. 10.1016/j.phrs.2018.09.006.

18. Sakai, S., Watanabe, S., Komine, O., Sobue, A., and Yamanaka, K. (2021). Novel reporters of mitochondria-associated membranes (MAM), MAMtrackers, demonstrate MAM disruption as a common pathological feature in amyotrophic lateral sclerosis. FASEB J 35, e21688. 10.1096/fj.202100137R.

19. Area-Gomez, E., de Groof, A.J., Boldogh, I., Bird, T.D., Gibson, G.E., Koehler, C.M., Yu, W.H., Duff, K.E., Yaffe, M.P., Pon, L.A., and Schon, E.A. (2009). Presenilins are enriched in endoplasmic reticulum membranes associated with mitochondria. Am J Pathol 175, 1810–1816. 10.2353/ajpath.2009.090219.

20. Naon, D., Zaninello, M., Giacomello, M., Varanita, T., Grespi, F., Lakshminaranayan, S., Serafini, A., Semenzato, M., Herkenne, S., Hernandez-Alvarez, M.I., et al. (2016). Critical reappraisal confirms that Mitofusin 2 is an endoplasmic reticulum-mitochondria tether. Proc Natl Acad Sci U S A 113, 11249–11254. 10.1073/pnas.1606786113.

21. Hirabayashi, Y., Kwon, S.K., Paek, H., Pernice, W.M., Paul, M.A., Lee, J., Erfani, P., Raczkowski, A., Petrey, D.S., Pon, L.A., and Polleux, F. (2017). ER-mitochondria tethering by PDZD8 regulates Ca(2+) dynamics in mammalian neurons. Science 358, 623–630. 10.1126/science.aan6009.

22. Hayashi, T., Rizzuto, R., Hajnoczky, G., and Su, T.P. (2009). MAM: more than just a housekeeper. Trends Cell Biol 19, 81–88. 10.1016/j.tcb.2008.12.002.

23. Pham, J.H., and Stankowska, D.L. (2023). Mitochondria-associated endoplasmic reticulum membranes (MAMs) and their role in glaucomatous retinal ganglion cell degeneration-a mini review. Front Neurosci 17, 1198343. 10.3389/fnins.2023.1198343.

24. Morcillo, P., Kabra, K., Velasco, K., Cordero, H., Jennings, S., Yun, T.D., Larrea, D., Akman, H.O., and Schon, E.A. (2024). Aberrant ER-mitochondria communication is a common pathomechanism in mitochondrial disease. Cell Death Dis 15, 405. 10.1038/s41419-024-06781-9.

25. Kannan, M., Lahiri, S., Liu, L.K., Choudhary, V., and Prinz, W.A. (2017). Phosphatidylserine synthesis at membrane contact sites promotes its transport out of the ER. J Lipid Res 58, 553–562. 10.1194/jlr.M072959.

26. Area-Gomez, E., Del Carmen Lara Castillo, M., Tambini, M.D., Guardia-Laguarta, C., de Groof, A.J., Madra, M., Ikenouchi, J., Umeda, M., Bird, T.D., Sturley, S.L., and Schon, E.A. (2012). Upregulated function of mitochondria-associated ER membranes in Alzheimer disease. EMBO J 31, 4106–4123. 10.1038/emboj.2012.202.

27. Larrea, D., Tamucci, K.A., Kabra, K., Velasco, K.R., Yun, T.D., Pera, M., Montesinos, J., Agrawal, R.R., Paradas, C., Smerdon, J.W., et al. (2025). Altered mitochondria-associated ER membrane (MAM) function shifts mitochondrial metabolism in amyotrophic lateral sclerosis (ALS). Nat Commun 16, 379. 10.1038/s41467-024-51578-1.

28. Johri, A., and Chandra, A. (2021). Connection Lost, MAM: Errors in ER-Mitochondria Connections in Neurodegenerative Diseases. Brain Sci 11. 10.3390/brainsci11111437.

29. Okumura, N., Kitahara, M., Okuda, H., Hashimoto, K., Ueda, E., Nakahara, M., Kinoshita, S., Young, R.D., Quantock, A.J., Tourtas, T., et al. (2017). Sustained Activation of the Unfolded Protein Response Induces Cell Death in Fuchs’ Endothelial Corneal Dystrophy. Invest Ophthalmol Vis Sci 58, 3697–3707. 10.1167/iovs.16-21023.

30. Onishi, T., Yuasa, T., Ueda, N., Miyadai, K., Tourtas, T., Schlotzer-Schrehardt, U., Kruse, F., Koizumi, N., and Okumura, N. (2025). The PERK-p38 MAPK Axis Drives Endoplasmic Reticulum Stress-Induced Apoptosis in Fuchs Endothelial Corneal Dystrophy. Invest Ophthalmol Vis Sci 66, 63. 10.1167/iovs.66.11.63.

31. Kumar, V., and Jurkunas, U.V. (2021). Mitochondrial Dysfunction and Mitophagy in Fuchs Endothelial Corneal Dystrophy. Cells 10. 10.3390/cells10081888.

32. Czarny, P., Seda, A., Wielgorski, M., Binczyk, E., Markiewicz, B., Kasprzak, E., Jimenez-Garcia, M.P., Grabska-Liberek, I., Pawlowska, E., Blasiak, J., et al. (2014). Mutagenesis of mitochondrial DNA in Fuchs endothelial corneal dystrophy. Mutat Res 760, 42–47. 10.1016/j.mrfmmm.2013.12.001.

33. Bannon, S.T., Shatz, N., Wong, R., Parekh, M., and Jurkunas, U.V. (2024). MitoQ relieves mitochondrial dysfunction in UVA and cigarette smoke-induced Fuchs endothelial corneal dystrophy. Exp Eye Res, 110056. 10.1016/j.exer.2024.110056.

34. Kasi, A., Steidl, W., and Kumar, V. (2025). Endoplasmic Reticulum-Mitochondria Crosstalk in Fuchs Endothelial Corneal Dystrophy: Current Status and Future Prospects. Int J Mol Sci 26. 10.3390/ijms26030894.

35. Qureshi, S., Lee, S., Ritzer, L., Kim, S.Y., Steidl, W., Krest, G.J., Kasi, A., and Kumar, V. (2024). ATF4 regulates mitochondrial dysfunction, mitophagy, and autophagy, contributing to corneal endothelial apoptosis under chronic ER stress in Fuchs’ dystrophy. bioRxiv. 10.1101/2024.11.14.623646.

36. Matusmoto, S., Kadoya, S., Horiuchi, Y., Okuda, H., Miyadai, K., Shima, Y., Young, R.D., Quantock, A.J., Schlotzer-Schrehardt, U., Kruse, F., et al. (2025). Enhanced mitochondria-associated membrane formation in Fuchs endothelial corneal dystrophy: a novel link between endoplasmic reticulum stress and mitochondrial dysfunction. Jpn J Ophthalmol. 10.1007/s10384-025-01288-y.

37. Guardia-Laguarta, C., Area-Gomez, E., Rub, C., Liu, Y., Magrane, J., Becker, D., Voos, W., Schon, E.A., and Przedborski, S. (2014). alpha-Synuclein is localized to mitochondria-associated ER membranes. J Neurosci 34, 249–259. 10.1523/JNEUROSCI.2507-13.2014.

38. Annunziata, I., Weesner, J.A., and d’Azzo, A. (2021). Isolation of Mitochondria-Associated ER Membranes (MAMs), Synaptic MAMs, and Glycosphingolipid Enriched Microdomains (GEMs) from Brain Tissues and Neuronal Cells. Methods Mol Biol 2277, 357–370. 10.1007/978-1-0716-1270-5_22.

39. Liu, Y., Ma, X., Fujioka, H., Liu, J., Chen, S., and Zhu, X. (2019). DJ-1 regulates the integrity and function of ER-mitochondria association through interaction with IP3R3-Grp75-VDAC1. Proc Natl Acad Sci U S A 116, 25322–25328. 10.1073/pnas.1906565116.

40. Filadi, R., Greotti, E., Turacchio, G., Luini, A., Pozzan, T., and Pizzo, P. (2015). Mitofusin 2 ablation increases endoplasmic reticulum-mitochondria coupling. Proc Natl Acad Sci U S A 112, E2174–2181. 10.1073/pnas.1504880112.

41. Hedskog, L., Pinho, C.M., Filadi, R., Ronnback, A., Hertwig, L., Wiehager, B., Larssen, P., Gellhaar, S., Sandebring, A., Westerlund, M., et al. (2013). Modulation of the endoplasmic reticulum-mitochondria interface in Alzheimer’s disease and related models. Proc Natl Acad Sci U S A 110, 7916–7921. 10.1073/pnas.1300677110.

42. Bravo, R., Vicencio, J.M., Parra, V., Troncoso, R., Munoz, J.P., Bui, M., Quiroga, C., Rodriguez, A.E., Verdejo, H.E., Ferreira, J., et al. (2011). Increased ER-mitochondrial coupling promotes mitochondrial respiration and bioenergetics during early phases of ER stress. J Cell Sci 124, 2143–2152. 10.1242/jcs.080762.

43. Miyai, T., Toyono, T., Kitamoto, K., Fukushima, M., Yoshida, J., Shirakawa, R., Nakagawa, S., Jurkunas, U.V., and Usui, T. (2018). Endoplasmic reticulum stress decreases mitochondrial membrane potential and upregulates PARK2 expression in corneal endothelium. Investigative Ophthalmology & Visual Science 59, 4436–4436.

44. Kim, M., Nikouee, A., Sun, Y., Zhang, Q.J., Liu, Z.P., and Zang, Q.S. (2022). Evaluation of Parkin in the Regulation of Myocardial Mitochondria-Associated Membranes and Cardiomyopathy During Endotoxemia. Front Cell Dev Biol 10, 796061. 10.3389/fcell.2022.796061.

45. Cali, T., Ottolini, D., Negro, A., and Brini, M. (2013). Enhanced parkin levels favor ER-mitochondria crosstalk and guarantee Ca(2+) transfer to sustain cell bioenergetics. Biochim Biophys Acta 1832, 495–508. 10.1016/j.bbadis.2013.01.004.

